# ESGI: Efficient splitting of generic indices in single-cell sequencing data

**DOI:** 10.64898/2026.03.04.709594

**Authors:** Tim Stohn, Nadine D. van de Brug, Anastasia Theodosiadou, Bram Thijssen, Katarzyna Jastrzebski, Lodewyk F.A. Wessels, Evert Bosdriesz

## Abstract

Single-cell sequencing technologies increasingly rely on complex nucleotide barcoding schemes to encode cellular identities, experimental conditions, and multiple molecular modalities within a single experiment. While demultiplexing, alignment, and UMI-based quantification form the core preprocessing steps that transform raw sequencing reads into analyzable single-cell data, existing pipelines are often tightly coupled to specific experimental designs and typically assume fixed barcode positions and substitution-only error models. As a result, many emerging assays employing combinatorial, variable-length, or multimodal barcoding designs require custom, hard-coded preprocessing solutions that are difficult to generalize and maintain.

Here, we present ESGI (Efficient Splitting of Generic Indices), a flexible and extendable framework for demultiplexing and processing single-cell sequencing data with arbitrary barcode architectures. ESGI operates directly on raw FASTQ files using a generic barcode pattern specification, supports barcode matching with insertions and deletions via Levenshtein distance, accommodates variable-length barcodes, and provides detailed quality metrics for barcode assignment. ESGI optionally integrates genome alignment via STAR and performs feature quantification and UMI collapsing to generate cellby-feature count matrices. ESGI is well documented and readily applicable to novel single-cell experiments.

We demonstrate the versatility of ESGI across six datasets spanning four distinct single-cell technologies, including combinatorial indexing–based transcriptomic and multimodal assays, feature barcode–based protein measurements, and spatial barcoding data. Across these applications, ESGI robustly demultiplexes complex barcode designs that are not natively supported by existing pipelines, while producing results comparable to established workflows where applicable. Together, ESGI provides a general and future-proof solution for preprocessing single-cell sequencing data, enabling rapid adoption and analysis of emerging experimental designs.

## 1 INTRODUCTION

Single-cell sequencing has evolved from scRNA-seq assays to a rapidly growing array of sequencing-based technologies that measure diverse molecular modalities at single-cell resolution. Central to many of these technologies is the use of nucleotide barcodes, which are employed to uniquely label individual cells, spatial coordinates [1], experimental conditions, modalities like transcriptome [2–4], protein [5, 6], chromatin accessibility [7], guides in CRISPR-based single-cell perturbations [8], combinations of them [9–11] and much more. The process of identifying the barcodes present in a sequencing read and thereby assigning the reads to their respective categories, such as individual cell identities or experimental conditions, is commonly referred to as demultiplexing. The measured features may comprise nucleotide sequences, such as DNA or RNA, which must be aligned to a reference genome prior to quantification, or feature barcodes, such as nucleotide-tagged antibodies whose sequences provide direct readouts of, for example, the abundance of the proteins bound by the antibodies. Single-cell identities are generally encoded either by a single barcode [11, 12] or by a combination of multiple barcodes [3, 6]. Following demultiplexing and mapping to the reference genome, the measured modalities are quantified for all cells. This is often done using unique molecular identifiers (UMIs) to identify PCR duplicates and obtain accurate molecule-level abundance estimates. Together, demultiplexing, mapping of genomic sequences and counting single-cell features (UMI collapsing) constitute the core processing steps that transform raw sequencing reads into cell-by-feature count matrices for downstream single-cell analysis. As barcoding schemes have increased in number and complexity, demultiplexing has become an increasingly technically challenging preprocessing step.

Several tools exist to process barcoding-based single-cell data, but most are designed around specific experimental platforms and offer limited flexibility for novel barcoding schemes. Commercial platforms such as 10x Genomics provide dedicated pipelines that are robust and well-maintained, for example Cell Ranger for 10x Genomics, but these are tightly coupled to their respective assay designs. STARsolo is a more flexible tool that performs integrated alignment, barcode processing, and gene quantification for a range of single-cell assays, and supports flexible barcode positioning within a single read, but cannot handle barcode segments spanning both reads of a paired-end library [13]. zUMIs supports user-defined barcoding schemes for single-cell RNA-seq data but is not designed for feature-barcodes or multimodal assays [14]. Alevin and its successor alevin-fry provide efficient, alignment-free quantification with flexible specification of barcode and UMI positions [15, 16]. kallisto and bustools enable rapid, pseudoalignment-based processing of single-cell data through a modular framework that accommodates arbitrary and combinatorial barcode schemes [17], and the kITE workflow extends this framework to feature barcode assays [18]. SDRRanger was developed for joint single-cell RNA and DNA sequencing with support for flexible barcode positioning [19]. Across these tools, a common limitation is that barcodes are extracted at fixed, predefined positions within the read, meaning the starting position of each barcode element is assumed rather than inferred from the data.

A second, related limitation concerns the correction of mismatches in barcodes. Barcode sequence similarity is commonly measured using either Hamming or Levenshtein distance. Hamming distance counts positional mismatches between equal-length sequences, allowing only for substitutions, whereas Levenshtein distance additionally accounts for insertions and deletions (indels). Most tools rely exclusively on Hamming distance for barcode matching, and therefore only account for nucleotide substitutions and not indels during barcode mapping. This distinction matters because indels occurring early in a read shift the expected positions of all downstream barcode elements, causing them to be misassigned even when the barcodes themselves are error-free. Such positional shifts are particularly consequential in combinatorial barcoding schemes, where multiple barcode elements are concatenated in a single read. These errors in barcodes can arise from various sources, such as errors in the synthesis of barcodes, errors during library preparation due to PCR amplification errors, or sequencing errors. Library preparation errors have been reported to occur at negligible rates below 10^−5^ per base for high-fidelity DNA polymerases [20]. Illumina short-read sequencing exhibits per-base error rates on the order of 10^−3^–10^−2^, with sequencing errors dominated by single-base substitutions rather than insertions or deletions [21, 22]. However, deletions are reported to be the most dominant errors in barcode synthesis [23], and Hawkins et al. explicitly show that deletions dominate synthesis errors in their barcoding experiments [24]. To address this, they recently developed an indel-correcting method called FREEbarcodes and implemented it in their demultiplexing pipeline SDRRanger [19], but indel-aware demultiplexing remains unavailable in most tools. Overall, current demultiplexing tools are frequently limited by fixed barcode extraction and substitution-only error correction. This prevents their application to complex barcoding patterns that demand both mapping flexibility and indel-aware barcode handling. To allow for flexible pattern matching *and* handle indels in barcodes, we developed ESGI (Efficient Splitting of Generic Indices), a flexible demultiplexing and processing framework designed to support complex single-cell barcoding schemes. ESGI takes as input raw FASTQ files together with a flexible barcode pattern specification that describes the order and allowed sequences of barcodes within each read. Based on this specification, ESGI performs barcode demultiplexing, optionally aligns genomic read sequences to a reference genome using STAR, and subsequently counts single-cell features and collapses UMIs to generate a final single-cell feature matrix. In contrast to existing tools, ESGI supports barcode matching with insertions and deletions via Levenshtein distance, can map variable-length barcodes (staggers) [2], and reports detailed quality metrics that provide insight into barcode assignment and mapping performance. As single-cell technologies continue to evolve toward increasingly complex barcoding patterns and multimodal designs, ESGI offers a flexible and easy-to-use solution for emerging methods that lack dedicated bioinformatics pipelines, while providing additional statistical information that may inform further optimization of experimental protocols. ESGI is extensively documented, and we provide vignettes to run ESGI on diverse single-cell technologies.

To demonstrate its versatility and performance, we applied ESGI to six datasets representing four distinct barcoding-based single-cell sequencing technologies. The datasets cover various barcoding technologies as well as RNA and protein measurements. The technologies include combinatorial indexing data profiling both RNA and protein abundance, a 10x-based technology measuring RNA and protein simultaneously and a novel spatial barcoding technology. First, to demonstrate the advantages of flexible and indel-aware debarcoding we applied ESGI to two combinatorial indexing datasets: data generated with SIGNAL-seq, a recently developed single-cell method employing combinatorial barcoding to jointly profile RNA and protein modalities [6], and a dataset generated using the SPLiT-seq technology [25], representing one of the original combinatorial barcoding approaches for single-cell transcriptomics. Next, we applied ESGI to Phospho-seq, which simultaneously quantifies RNA and protein abundances using 10x Genomics single-cell barcoding in combination with custom nucleotide-tagged antibodies [26]. Finally, we evaluated ESGI on xDBiT data, a parallelized spatial barcoding technology that employs a custom barcode design to encode spatial locations together with sample indices to process nine samples in parallel [1]. While the original xDBiT workflow relies on a custom pipeline built upon tools developed for Drop-seq data [27], we show that ESGI can robustly process this dataset with its flexible framework.

## 2 METHODS

### 2.1 Overview of ESGI

ESGI enables flexible demultiplexing and counting of barcoding-based single-cell data, such as multimodal, combinatorial barcoding or spatial data (Fig. 1a). Input to ESGI are the FASTQ files that have to be processed together with a user-provided barcode pattern (Fig. 1b). The predefined generic input pattern allows for various elements at different positions in the FASTQ reads, so ESGI can easily be applied to novel sequencing technologies with new barcoding combinations. This input pattern consists of several elements that together give a generic description of the expected nucleotide sequence in the FASTQ reads. These elements can be barcode elements where a sequence from a whitelist of potential barcodes can occur, constant elements, unique molecular identifiers (UMI) or genomic sequences like DNA/RNA. Barcode elements in the pattern can have many possible barcode sequences that can, for example, encode single-cell indices, feature indices or hash tags. Constant elements are predefined nucleotide sequences that, for example, link other elements in the pattern. Additionally, the user can set the number of allowed mismatches for every element individually or decide if a barcode should be mapped by Levenshtein distance, which allows for both substitutions and indels, or only by Hamming distance, which allows for substitutions only, but can be more efficient in some instances.

**Figure 1:**
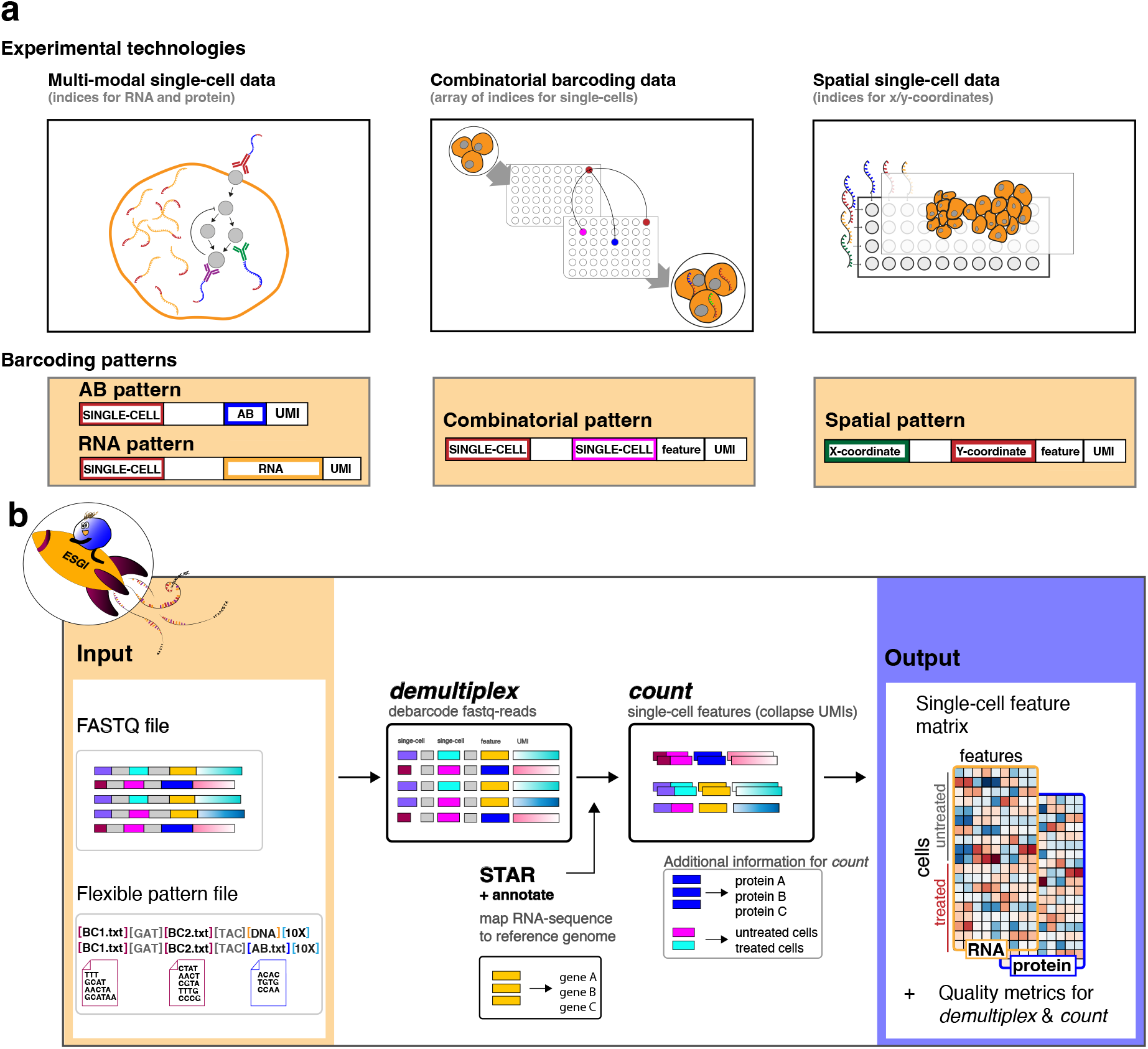
Workflow of ESGI. (a) ESGI can demultiplex various data types: multimodal data, with an RNA-sequence and an antibody-attached (AB) feature barcode encoding the protein modality, combinatorial barcoding data, where the combination of barcodes encodes for individual cells, or spatial data with barcodes encoding coordinates. (b) Input to ESGI are the raw FASTQ files and a flexible pattern which describes the barcoding scheme as a list of barcode elements. ESGI then maps the patterns to the FASTQ reads. ESGI is comprised of two main tools: `demultiplex` and `count`. `demultiplex` maps the expected barcodes in the FASTQ file and `count` counts the number of reads for single-cell features by collapsing UMIs. If the data contains a ‘DNA’ element, ESGI can call STAR and create a genome annotation that is added to the output of `demultiplex` with the tool `annotate` before calling count.

ESGI can be split into two separate steps: demultiplexing the sequence pattern and counting singlecell features. ESGI can be run as a wrapper of these two tools, but it can also be split into separate calls of the internal tools `demultiplex` and `count` for more control. During the demultiplexing step the FASTQ reads are split into the elements as defined in the input pattern. Elements are corrected if they are within the mismatch limit. If genomic sequences are present, ESGI can call STAR to map these sequences to a reference genome and add the STAR annotation to the output of `demultiplex`. The counting step collapses UMIs, generates a single-cell feature matrix, and assigns additional barcoded attributes, like treatment conditions, if these are present in the barcode pattern. In addition to the final data matrix ESGI generates quality metrics for the demultiplexing and counting step.

### 2.2 Demultiplexing

ESGI demultiplexes barcoding-based single-cell data by mapping the input reads to a predefined generic input pattern. In this pattern, every barcode element is enclosed within square brackets and can consist of constant linker sequences, barcode elements, random elements or a genomic sequence. Constant linker elements are described by the exact nucleotide sequence. Barcode elements, which can, for example, be used to identify individual cells, are specified in whitelist files (e.g., ‘BC1.txt’) that contain all valid barcode sequences for this barcode element. ‘DNA’ denotes genomic or transcriptomic sequence content to be aligned to a reference genome, and random sequences of length *L*, such as UMIs, are described as ‘*LX*’ (Fig. 2b). ESGI maps individual pattern elements sequentially along the read allowing for substitutions and indels. To do so, ESGI precomputes all possible sequences with a Hamming distance of one for each constant element and all barcodes of the barcode elements. ESGI starts with the first pattern element, extracts the expected sequence length from the input read, and checks whether the extracted sequence matches any of the expected barcodes or the precomputed sequences with a mismatch. By default, ESGI also allows for insertions and deletions. To that end, ESGI also precomputes all possible sequences with one insertion or deletion (unless the user explicitly requests Hamming distance only). The start of the following pattern element is defined by the end position of the currently mapped pattern element. In contrast to extracting pattern elements from fixed positions, this approach prevents shifts in the mapping frame that would otherwise propagate to downstream barcodes (Fig. 2a).

**Figure 2:**
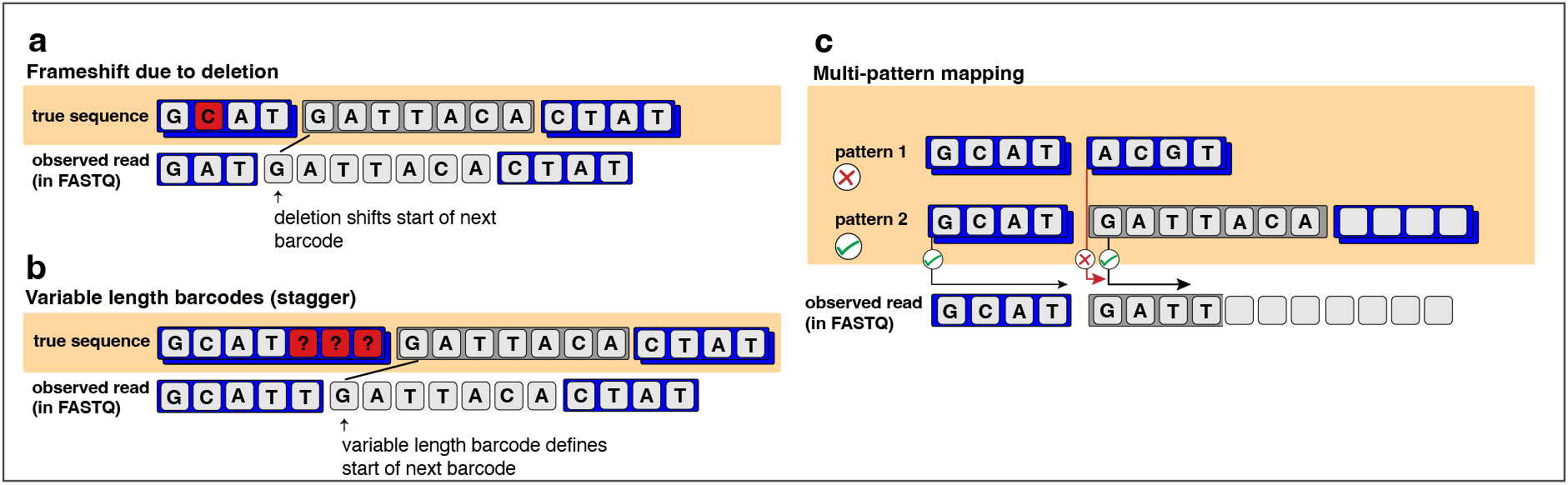
ESGI’s flexible framework enables effective handling of novel and previously unaddressed scenarios. (a) ESGI maps barcodes allowing for substitutions and indels. Indels can lead to a shift in the mapping-frame which results in new starting positions for subsequent barcodes. (b) Variable length barcodes can make it impossible to ‘cut’ barcodes at fixed positions in the read. By mapping barcodes sequentially ESGI can map barcodes of variable length and starts at the new offset with the subsequent barcode. Therefore, the starting positions of barcodes depends on the mapping of the previous barcode. (c) ESGI can demultiplex several patterns at once in cases where a FASTQ file contains several modalities that were captured and sequenced together.

If a pattern contains barcode elements and more than one mismatch is allowed, ESGI aligns potential barcode sequences to the input reads. For barcode elements ESGI uses positional k-mers to reduce the search space for possible barcodes at this position. To do so, ESGI calculates all positional k-mers of the whitelist barcodes and before mapping filters out barcodes that do not share a minimum number of positional k-mers with the extracted sequence (Fig. S2). The remaining barcodes are then aligned to the input sequence using a bit-parallel algorithm as implemented in the Edlib library [28]. ESGI aligns barcodes to the input reads using a semi-global alignment strategy. This approach permits unpenalized terminal deletions in the extracted sequence that is mapped to the barcode, and thereby prevents excess bases at the end of the extracted sequence that do not map to the barcode from being counted (for instance when a deletion in the barcode sequence shortens the effective barcode length in the extracted sequence) (Fig. S3). As a result mismatches are not artificially inflated, in contrast to a pure Levenshtein distance computed between equal-length sequences, similar to the idea of FREE-barcodes [24]. To further improve the running time, ESGI precomputes all pairwise distances between barcodes of a barcode element if the number of possible barcodes is below 500. During mapping, these distances are used to terminate alignments early if all remaining unchecked barcodes must exceed the current best alignment score. The precomputed distances yield a matrix of shortest possible edit distances between all theoretically attainable barcodes. During the alignment of barcodes to the FASTQ reads, this information allows ESGI to terminate the search early once the closest possible match is identified. Concretely, suppose a candidate barcode *b* aligns to a sequence *s* with edit distance *d* (*b, s*). Let *d*_min_(*b, b*^′^) denote the minimum edit distance between *b* and any other barcode *b*^′^ ≠ *b*. If *d* (*b, s*) *< d*_min_(*b, b*^′^) − *d* (*b; s*), then no alternative barcode can achieve a smaller alignment distance to the sequence. Hence, *b* is guaranteed to be the optimal match, and ESGI can safely terminate the alignment procedure. ESGI discards reads if the detected barcode at a barcode element is ambiguous and more than one barcode would map with a minimal edit distance.

Barcoding-based single-cell data can contain barcodes that are of variable length. These are often introduced by staggers, which result in offsets in barcodes and are useful to increase base diversity and thereby improve cluster detection and base calling during sequencing [29]. However, variable length barcodes cannot be ‘cut out’ at fixed positions in the sequence. Therefore, ESGI tries to map all possible barcodes with variable lengths, reports the result if one single barcode fits best with minimal edit-distance and continues mapping the subsequent pattern element at the end position of this sequence (Fig. 2b).

ESGI can simultaneously map multiple patterns to a FASTQ file, enabling demultiplexing when sequencing libraries have not been separated prior to sequencing (Fig. 2c). ESGI therefore tries to map all possible patterns, and if one pattern matches with minimal edit-distance reports this pattern. This capability also allows the specification of more flexible barcode schemes, for example, when the validity of one pattern element depends on the identity of another element and only specific barcode combinations are permitted. By supporting multiple patterns, ESGI simplifies the demultiplexing process and makes conditional and hierarchical barcode designs possible (Supplementary Information A.1).

### 2.3 Calling STAR

ESGI calls STAR (version ≥ 2.7.9) as an external aligner to obtain genomic coordinates and gene annotations. `demultiplex` separates the genomic region of the input reads from the remaining pattern elements and writes the genomic reads into a FASTQ file and the remaining pattern elements into a.tsv file. STAR is invoked in non-solo mode to produce genomic BAM files containing GX and GN tags for the FASTQ file containing only the genomic reads without barcoding information, where GX denotes the Ensembl gene identifier (ENSG) and GN the corresponding gene symbol. ESGI then extracts these tags via an internal annotation step and associates each read with its gene identity. The resulting gene annotations are merged with the demultiplexed pattern elements (.tsv file) to generate demultiplexed reads with gene-annotations, which can be used as input to `count`.

### 2.4 Feature counting

The final step of ESGI is counting single-cell feature occurrences with the tool `count`. Input for this step is either the output of `demultiplex` directly or the STAR-annotated output of `demultiplex` if genomic sequences (as in scRNA-seq) are present. In this step UMIs are collapsed and counted to generate the final single-cell feature matrix. First, reads are sorted by single-cell and feature barcodes and UMIs are only collapsed if they come from the same feature in the same cell. Therefore, UMIs are initially counted based on exact matches and subsequently ordered by their count. For the same single-cell feature combination identical UMIs are collapsed into a single count for that feature in that cell. Sequencing errors can result in UMIs originally arising from the same molecule to differ in the sequenced read. To account for this, we employ a strategy similar to alevin [15]. Specifically, starting with the most abundant UMI, we compare it to all UMIs from the same feature in the same single-cell with equal or fewer counts. If the distance between two UMIs is below a threshold, the less abundant UMI can be collapsed into the more abundant one (by default allowing a Hamming distance of one). UMIs are only collapsed if the less abundant UMI has less than 20% of the counts of the more abundant UMI. For more detailed information see Supplementary Information A.3.

### 2.5 Benchmarking ESGI on SIGNAL-seq data

To demonstrate the applicability of ESGI to novel and highly multiplexed barcoding schemes, we applied ESGI to a dataset generated using SIGNAL-seq technology, which labels individual cells through three rounds of combinatorial barcoding and jointly profiles RNA and protein modalities using feature barcodes [6]. We used the datasets of HeLa spheroid cells which are publicly available from GEO (accession GSE256405), which were downloaded from the Sequence Read Archive (SRA run SRR28056729 for the RNA modality and SRA run SRR28056728 for the protein modality).

The authors of SIGNAL-seq processed the protein modality with kITE [18]. Since kITE only maps barcodes at the expected positions we ran ESGI without explicitly mapping the constant linker regions between barcode elements unless stated differently. As kITE permits one substitution per barcode element, we applied the same mismatch tolerance in ESGI by allowing one mismatch in each barcode element. kITE does not allow for mismatches in the extracted UMI. Therefore, we also ran ESGI with zero mismatches in the UMI during counting of the single-cell feature barcodes with `count`. The authors of SIGNAL-seq processed the RNA modality with zUMIs [14]. zUMIs concatenates the three barcode sequences before error correction, and we ran it with three substitutions for the concatenated sequence. Similarly, we allowed 1 mismatch in every barcode element with ESGI. For the RNA modality, both zU-MIs and ESGI were executed with UMI error correction permitting a single nucleotide substitution in the UMI sequence. Since ESGI supports only counting of exonic reads we ran zUMIs without counting intronic reads. The data contains reads from oligo-dT and random hexamer reverse transcription primers and both reads have different barcodes. zUMIs was run to assign both reads to the same cell and we therefore also ran ESGI by supplying a barcode-mapping file, which combines barcodes and accounts reads from both barcodes to the same cell. To compare the results obtained by ESGI to those obtained by kITE/zUMIs, we computed per-cell correlations between feature counts of the compared tools across all measured features. For kITE we took the unfiltered counts and removed cells for which correlations could not be computed (due to no detected features or zero variance). For the comparison of RNA counts we separately prefiltered the output of ESGI and zUMIs to include only cells for which the tools detected more than 100 counts, calculated the per-cell correlations and also excluded cells for which correlations could not be computed.

### 2.6 Benchmarking ESGI on Phospho-seq data

Many single-cell experiments are run with 10x or are a combination of custom protocols that are combined with 10x. While 10x data can be easily demultiplexed and processed with Cell Ranger, custom protocols might need adjustment. Therefore, to show ESGI’s compatibility with 10x and its ability to build upon their barcoding scheme, we processed scRNA-seq and protein quantification data of the Phospho-seq technology with ESGI [26]. This dataset contains a single 10x single-cell barcode, while the protein modality is encoded using feature barcodes. We downloaded two datasets of the Phospho-seq-Multi experiment for the RNA and protein modality. The data is publicly available from GEO (accession GSE285561) and was downloaded from the Sequence Read Archive (SRA run SRR31955816 for the RNA modality and SRR31955815 for the protein modality). The authors processed the protein modality with alevin and the RNA modality with 10x Genomics Cell Ranger. Therefore, we processed both modalities in a similar fashion with ESGI and compared the results to the results of alevin for the protein and with Cell Ranger for the RNA modality. For the comparison of ESGI with Cell Ranger we generated a GRCh38 reference using Cell Ranger (v9.0.1) with Gencode v43. The resulting Cell Ranger–processed genome FASTA and gene annotation were reused to construct a separate STAR (v2.7.10b) genome index for the ESGI pipeline, ensuring identical reference sequences and gene models across analyses. We excluded intronic regions for the Cell Ranger run for comparability with ESGI. We then ran ESGI allowing for one mismatch in the single-cell barcode and collapsed UMIs with a Hamming distance of one. To calculate correlations between the results obtained by the methods, we used the raw unfiltered output of Cell Ranger and excluded cells for which we could not calculate a correlation due to no detected features in one of the tools or zero variance. Alevin allows for one mismatch (Hamming distance) in the single-cell and feature barcode and we ran ESGI as well with one mismatch. Similar to alevin we ran ESGI allowing for one substitution in UMIs during UMI collapsing. To calculate correlations for the results of both tools we filtered single cells that had at least five counts across all measured proteins and we excluded cells that were not detected in one technology or had zero variance in the protein counts.

### 2.7 Benchmarking ESGI on xDBiT data

We also applied ESGI to spatial data. To showcase the ease of application to novel sequencing technologies we applied ESGI to a new multiplexed spatial technology, which barcodes spatial coordinates as well as sample identities to process nine tissue samples in parallel [1]. The data contains three barcode elements – one for each of the x and y-coordinates, as well as a third barcode to distinguish between nine tissue samples. The three elements are connected by two 30 nucleotide long constant linker elements. We analyzed the first replicate of the mouse Cerebellum tissue, which is publicly available from GEO (accession GSE207843) and was downloaded from the Sequence Read Archive (SRA run SRX16111595). We compared the output of ESGI to the output of the xDBiT toolbox using the precomputed single-cell count matrix provided by the authors via their GitHub repository ([xDBiT toolbox](https://github.com/jwrth/xDBiT_toolbox/tree/main)). We ran ESGI allowing for one mismatch in the barcode elements and when running ESGI in the mode where we also explicitly map the constant elements, we allowed 10 mismatches in both linker elements. We collapsed UMIs allowing for 1 substitution in the UMI sequence. For the correlation analysis we excluded cells for which no correlation could be computed due to missing counts or zero variance.

### 2.8 Analysis of SPLiT-seq data

SPLiT-seq (Split Pool Ligation-based Transcriptome sequencing) is a combinatorial indexing method for single-cell RNA sequencing that labels transcripts through successive rounds of split–pool barcoding [25]. In each round, cells are redistributed across wells and tagged with a new barcode, and the combination of barcodes uniquely identifies the cell of origin after sequencing. To count the number of mismatches in the SPLiT-seq data we ran `demultiplex` on a small mouse brain dataset of 100 nuclei that was generated with the SPLiT-seq technology and also used in a benchmarking study of pipelines for SPLiT-seq data [30]. The data contains three barcode elements that are eight nucleotides long and are connected by two 30 nucleotide long constant linker elements. For the demultiplexing we allowed one mismatch in the barcode elements and when mapping with the constant elements we permitted 10 mismatches in both. To count the number of mismatches we ran ESGI to extract only the three barcode elements that serve as single-cell indices and count the edit operations during mapping. The data is publicly available from GEO (accession GSE110823) and was downloaded from the Sequence Read Archive (SRA run SRR6750041).

## RESULTS

### 3.1 ESGI enables flexible pattern mapping

Most existing tools for barcoding-based single-cell sequencing data provide limited flexibility regarding barcode architecture, disallow insertions and deletions (indels) within barcode sequences, and lack support for mapping constant linker elements containing indels that might shift subsequent barcode positions. To address this, we developed ESGI to facilitate easy demultiplexing and counting even for complex barcoding technologies (Fig. 2). To showcase the usability of ESGI we selected six datasets from four novel technologies with custom barcoding schemes. We processed the data with ESGI and compared the output of ESGI with tools that were used to process the various datasets in the respective publications and all allow the mapping of a generic barcode pattern (Table 1). None of these pipelines supports indels or barcodes of variable length (staggers), because they all extract barcodes at predefined positions within the input read, whereas ESGI maps barcodes sequentially. Of note, alevin-fry does allow for variable-length barcodes for specific single-cell protocols such as inDropV2 and sci-RNA-seq3, but not for variable-length barcodes at any position within the read. Additionally, ESGI can process multiple barcode patterns simultaneously, enabling analysis of sequencing libraries that contain different modalities with distinct read structures or barcoding schemes with hierarchical organization and mutually-exclusive barcode sets (A.1).

**Table 1:**
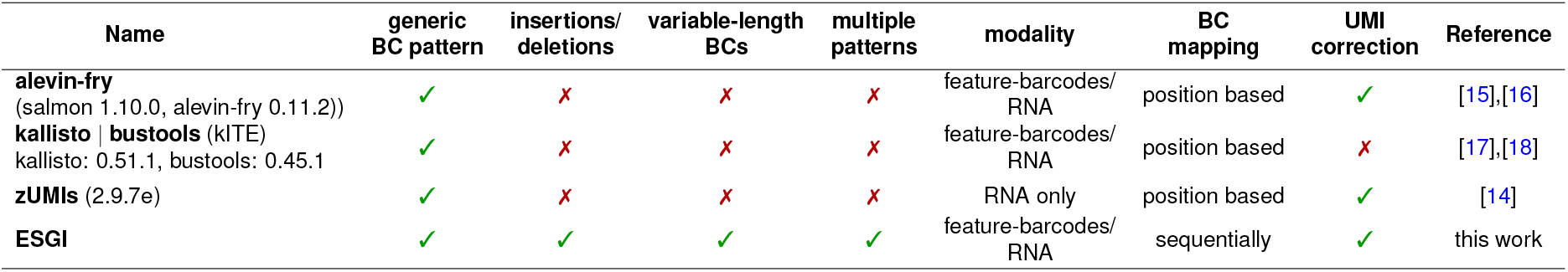
Comparison of demultiplexing tools or workflows for single-cell data used for benchmarking. The information shown is from installed tool versions as of December 2025. From the kallisto and bustools workflow we applied the kITE workflow for processing feature-barcode data. The table focuses on the flexibility of the pipelines with respect to processing diverse barcoding-based single-cell sequencing data types. We show whether the tools can process flexible barcode patterns (user defined barcode positions), can handle indels, barcodes of variable length at the same position or multiple patterns within one FASTQ file. We further consider the modalities the tools were developed for (feature barcodes or scRNA-seq data), whether they expect barcodes at predefined positions or map barcodes sequentially (the starting positions of a pattern element depends on the end position of the previous one) and if they perform UMI correction.

### 3.2 Indel-aware mapping improves read recovery

Combinatorial barcoding schemes rely on multiple barcode elements to encode single-cell identities, making them inherently sensitive to sequencing errors that occur early in the read. In particular, insertions or deletions shift downstream barcode positions and thereby compromise the accurate assignment of subsequent barcodes. To assess ESGI’s ability to robustly handle such error-prone read structures, we applied it to the multimodal SIGNAL-seq dataset [6], which profiles RNA and protein expression using three rounds of combinatorial barcoding. We demultiplexed both modalities using ESGI and compared the results to the original processing strategy, in which RNA reads were demultiplexed using zUMIs and protein-derived feature barcodes were processed using kITE, which relies on barcode matching with Hamming distance (Methods).

To evaluate to which extent indels occur and affect the mapping quality of barcodes in combinatorial barcoding experiments, we started by running ESGI on the RNA and protein modality and allowing for one mismatch (Levenshtein distance) in each of the three single-cell barcode sequences BC1, BC2 and BC3 (Fig. S5a). To enable comparison of the results with existing tools that often merely ‘cut out’ barcodes at the expected position in the sequence, we ran ESGI in a mode where we only mapped the barcodes and ignored the constant linker sequences between barcode elements since existing tools often do not take these constant elements into account. Since ESGI records the number of edit operations for each pattern element, a direct comparison of the frequencies of substitutions, insertions, and deletions can be made. The protein modality has almost as many deletions as substitutions, while the RNA modality has even more deletions than substitutions, and has an overall higher mismatch rate (Fig. 3a). We performed a similar analysis in an xDBiT and SPLiT-seq dataset. The barcoding pattern in both datasets comprises three barcode elements that are connected with constant linkers of 30 bases. In both datasets, deletions are almost as abundant or even more abundant than substitutions (Fig. S6a). This is in line with the reported higher deletion rates during barcode synthesis [23, 24], and highlights the need for indel-aware demultiplexing.

**Figure 3:**
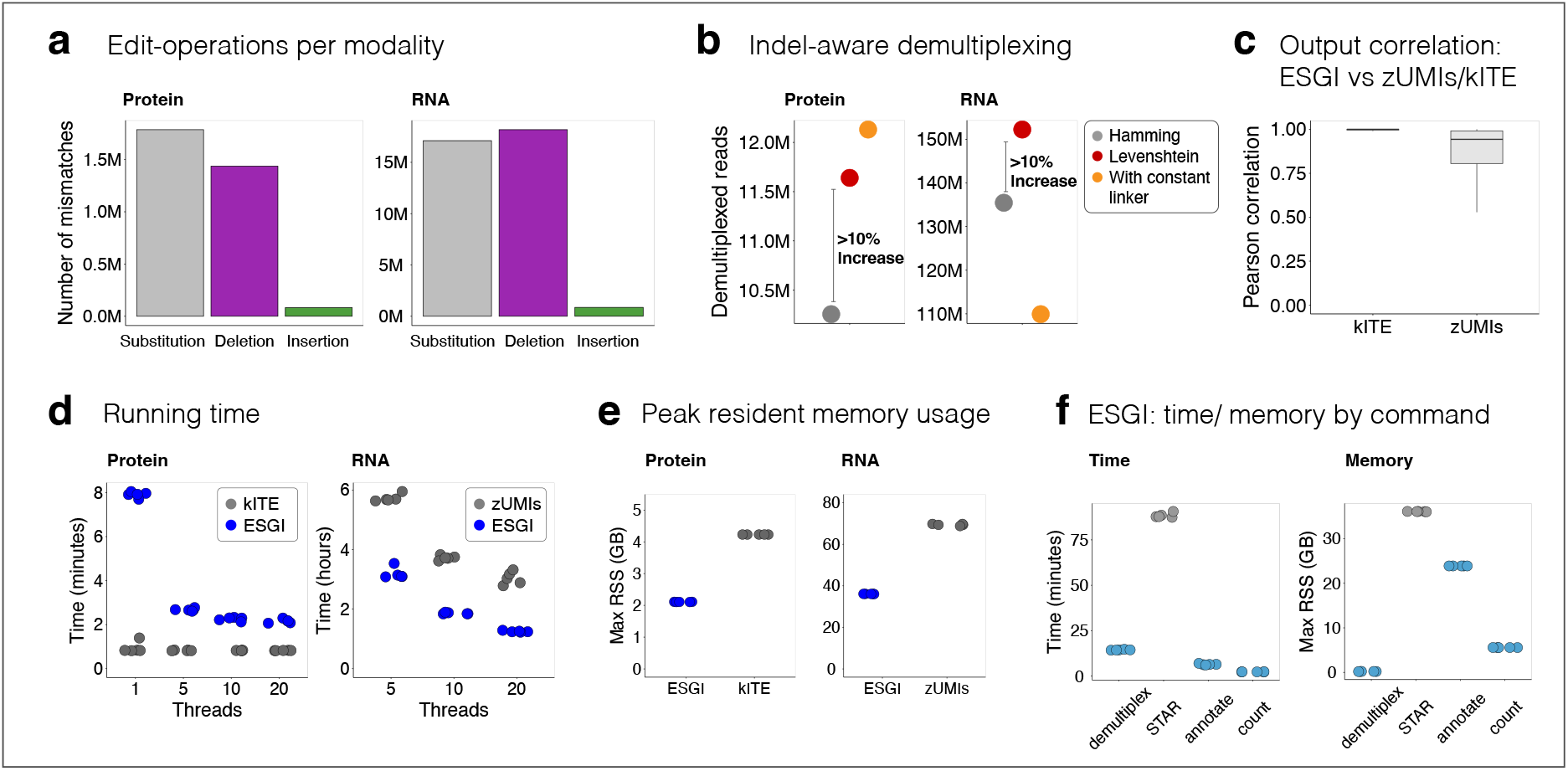
Application of ESGI to SIGNAL-seq data. (a) The number of edit-operations in the data of the protein and RNA modalities in the three *single-cell* barcode elements (BC1,BC2 and BC3) reported by ESGI when mapping only the three barcodes at fixed positions with one mismatch in each barcode (structure diagram (Fig. S5a)). (b) The number of successfully demultiplexed reads using ESGI under different matching strategies. Reads were demultiplexed allowing either a single substitution only (Hamming distance) or allowing a single substitution, insertion or deletion (Levenshtein distance), while mapping only the three barcode elements. In addition, ESGI was run with explicit mapping of the two constant elements. We permitted up to five mismatches for the protein modality in both constant elements and permitted up to 15 and 25 mismatches in the RNA modality, in the constant elements 1 and 2 respectively. (c) Pearson correlations between feature counts over all cells. (d) Running times of kITE, zUMIs and ESGI for the protein and RNA modality demultiplexing. All tools were run five times. (e) Peak resident memory usage for kITE, zUMIs and ESGI for the runs with ten threads. (f) Running time spent in the individual commands of ESGI as well as peak resident memory usage per command for ESGI during processing of the RNA modality with ten threads.

To quantify the benefit of allowing for indels, we demultiplexed both modalities in two ways: (1) allowing only for a Hamming distance of one in every barcode element, and (2) allowing for a Levenshtein distance of one. Allowing for indels increased the number of processed reads by more than 10% (Fig. 3b) for both modalities. The reads of the RNA modality had lower sequencing quality overall, so mismatches in the preceding constant linker elements could change the reading frame and thus lead to more mismatches in subsequent barcode elements (Fig. S5a). Therefore, we ran ESGI explicitly mapping the constant linker sequences between barcode elements, allowing for up to five mismatches (Levenshtein distance) in the constant elements. For the protein modality, this further improved the number of successfully processed reads. However, for the RNA modality, the number of successfully demultiplexed reads remains lower than when ESGI is run without the constant elements (even allowing for 15 and 25 mismatches in the two constant regions of length 22 and 30 respectively) (Fig. S5a). This is likely because the RNA modality had very low average base quality, especially in the second constant linker sequence. Many reads contained more than ten mismatches, which severely compromised reliable sequence mapping and led to a substantial reduction in the number of mapped reads (Fig. S5b). Allowing for indels also significantly increased the number of demultiplexed reads in the SPLiT-seq and xDBiT datasets, by more than 10% and 15%, respectively. Explicitly mapping the constant linker elements increased the number of demultiplexed reads only marginally (Fig. S6b). The Phospho-seq data on the other hand, which contains only a single round of barcoding plus an additional feature-barcode, contains mainly substitutions (Fig. S5a). For the protein modality, 41% of the reads matched the input pattern perfectly, and mapping with Hamming or Levenshtein distance improved the number of recovered reads only marginally (44% when mapping with Hamming distance and 45% with Levenshtein distance). Alevin shows a similarly low recovery rate with 42% of uniquely mapped reads. For the RNA modality, ESGI mapped 75% of the reads perfectly without mismatches and only an additional 1% could be mapped with 1 mismatch (Hamming) in the barcode element. To further compare the results of ESGI to the results of kITE and zUMIs, we generated single-cell count matrices from the demultiplexed reads for each tool, and computed per-cell correlations of ESGI to those obtained by kITE and zUMIs. The correlation of feature counts per cell is overall very high with a median of one for the comparison of kITE and ESGI and a median of 0.94 for zUMIs and ESGI (Fig. 3c). Of note, the increase in demultiplexed reads due to indel-aware mapping directly translated to an increase in detected single-cell feature counts for the protein data when comparing ESGI to kITE (Fig. S7a) and reads per cell for the RNA modality when comparing ESGI to zUMIs (Fig. S7b). Similarly, the xDBiT and Phospho-seq count matrices generated by ESGI correlate well with the results of the xDBiT-toolbox, alevin, and Cell Ranger (Fig. S6c). Together, these results indicate that indel-aware mapping increases the number of reads that can be assigned to a feature in a meaningful way.

### 3.3 ESGI enables efficient mapping with low memory usage and moderate running time

To compare the runtime and memory usage of ESGI with that of kITE and zUMIs, we ran the tools 5 times on the protein or RNA modalities of the SIGNAL-seq dataset. kITE processed the 17 million reads of the protein data in under one minute, while ESGI needed eight minutes when it was run with a single thread and about two minutes when run on 20 threads (Fig. 3d). For the RNA modality, ESGI processed the 183 million reads much faster than zUMIs, in about one hour compared to around three hours for zUMIs (Fig. 3d). Of note, ESGI spent more than 79% of the time running STAR. The peak memory usage of ESGI was around 2 GB for the protein and below 40 GB for the RNA dataset (Fig. 3e). This was almost two times lower than kITE or zUMIs. For the RNA data, the majority of the memory was again attributed to STAR and the annotation of the demultiplexed reads (Fig. 3f). ESGI can run `demultiplex` to map barcodes with either a Hamming distance or Levenshtein distance. When barcodes are precomputed, `demultiplex` runs in both cases at just below 50 seconds for the protein modality. Even when only precomputing barcodes with substitutions and aligning barcodes in cases when no valid barcode with substitutions was found, ESGI runs in less than 60 seconds (Fig. S7c).

### 3.4 ESGI provides additional information on the barcode mapping quality

To facilitate data quality control (QC), ESGI outputs position resolved information about the mapping quality of the barcoding pattern. If a read does not map to the predefined pattern, ESGI reports the pattern element where mapping failed. To showcase this functionality, we evaluated the quality metrics of the protein modality of the SIGNAL-seq data. SIGNAL-seq uses feature barcodes as readouts of protein abundances in single-cells and marks cells with a combination of three barcodes (Fig. 4a). Figure 4b shows at which position in the predefined input pattern the mapping failed when either allowing for zero mismatches or one mismatch in each of the barcode elements, and five in each of the constant elements. When allowing for zero mismatches in any pattern element, the mapping fails most often in the first, and only, element of the forward read and the first element of the reverse read (BC3). However, when allowing for mismatches, mapping fails more evenly across the pattern elements, with most errors occurring in the last position of the reverse read (BC1). Additionally, ESGI reports the number of mapped reads with and without mismatches for all positions in the pattern. Figure 4c shows the number of reads with zero or one mismatch in the three barcode elements. The difference between reads with zero and one mismatch progressively decreases from BC3 to BC1. As BC3 is positioned at the start of the read—where sequencing quality is generally highest—this may facilitate more accurate mapping and reduce the impact of mismatches. Some barcodes were detected more frequently with one mismatch than with zero mismatches in BC1 and the variance in the number of reads is much higher in BC1. Figure 4d shows the number of substitutions, deletions and insertions for the various pattern elements in the reverse read. Overall, the pattern elements show a high number of deletions, consistent with previously reported high deletion rates in synthetic barcodes [23, 24]. Especially the first constant region shows a high deletion rate. If pattern elements were not mapped sequentially these deletions could lead to a shift in the starting positions for pattern elements and barcodes could be missed.

**Figure 4:**
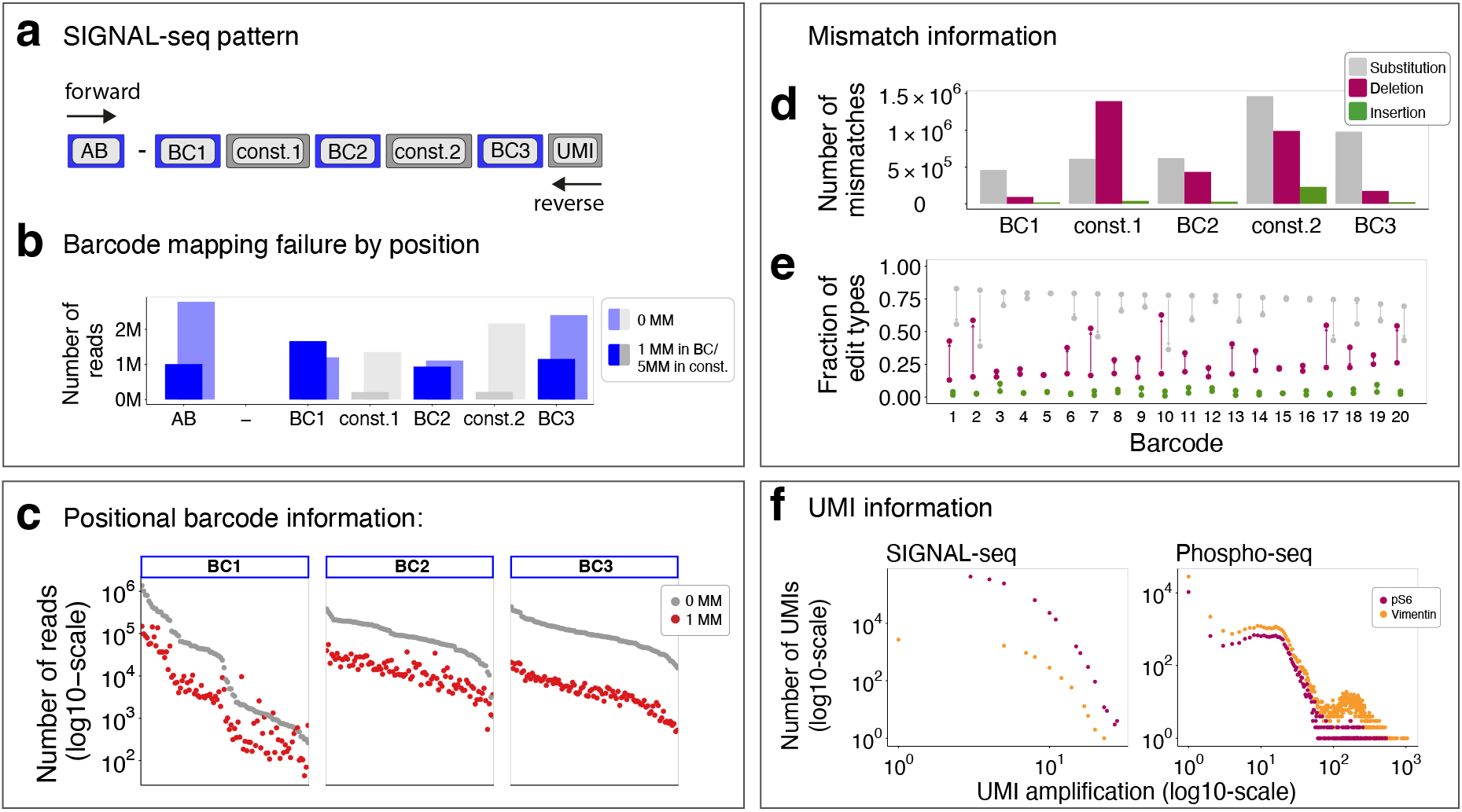
ESGI reports QC information. For the statistical information we explicitly mapped the whole pattern of the SIGNAL-seq data: the antibody-tagged feature barcode (AB), three barcode elements encoding the single-cell identity (BC) and also the constant elements (const.1 and const.2). (a) The input pattern describing the barcoding layout of the SIGNAL-seq data. (b) The number of reads that failed at the different positions of the input pattern. ESGI was run allowing for zero mismatches or for one mismatch in the barcode elements and five mismatches in the constant elements. (c) Number of successfully demultiplexed reads for all barcodes in all three barcode positions. Every dot represent one barcode and barcodes are ordered in decreasing order by the number of reads that contain this barcode with zero mismatches. ESGI was run allowing for at most one mismatch in every barcode element. Grey depicts the number of reads that have zero mismatches in a barcode and red shows the number of reads with one mismatch in a barcode. (d) Number of edit-operations (substitution, insertion, deletion) by position in the input pattern in the reads that were mapped successfully. (e) Fraction of reads with one substitution, insertion or deletion for the first 20 barcodes in alphabetical order of the barcode elements BC2 and BC3. Both barcode elements share the same barcodes and arrows connect the barcodes of BC3 with the barcodes of BC2. (f) Distribution of UMI amplification for the two most abundant feature barcodes in the SIGNAL-seq and Phospho-seq data respectively.

Additionally, ESGI stores the number of different edits for all possible barcodes in barcode elements. Figure 4e shows the fraction of reads with different edits for the barcodes in the barcode elements BC2 and BC3. The fraction of deletions goes up drastically in BC2, while the fraction of insertions remains stable. Some barcodes (e.g., barcode 2 and 10) show a marked increase in the fraction of deletions between barcode elements BC3 and BC2, whereas others (e.g., barcode 3 and 4) display little to no change in deletion frequency. These barcode-specific differences in the distribution of insertions, deletions, and substitutions provide insight into how individual barcodes are differentially affected by editing events. Such information may help identify barcodes that are more susceptible to certain types of sequence alterations and can therefore inform future barcode design and optimization strategies. Finally, ESGI counts the number of reads for every observed UMI (before UMI correction). This makes it possible to plot the frequency of UMI amplifications (Fig. 4f), which might serve as an additional metric to evaluate the sequencing depth or to filter out true molecule reads from background noise that might be introduced during PCR [31, 32]. We envision that the QC metrics as shown in Figure 4 provide a practical framework to improve or debug novel experimental technologies during their development phase and enable quantitative assessment of sequencing depth sufficiency.

## DISCUSSION

Barcoding-based sequencing protocols are rapidly increasing in complexity, incorporating multiple molecular modalities and increasingly elaborate barcoding schemes. In particular, combinatorial designs that rely on multiple rounds of barcoding can be affected by indels as insertions or deletions occurring early in the read can shift downstream barcode positions and substantially degrade mapping accuracy. ESGI addresses these challenges by providing a flexible and generic demultiplexing framework that supports barcodes of variable length at arbitrary positions within single- or paired-end reads and enables indel-aware barcode matching. Moreover, ESGI allows multiple barcode patterns present within a single FASTQ file to be processed simultaneously, facilitating hierarchical barcode assignment and the joint processing of multimodal libraries. Together, ESGI provides a flexible framework for demultiplexing diverse experimental designs and makes it well-suited for emerging novel sequencing technologies. ESGI makes it possible to quickly go from raw FASTQ files to the final single-cell feature matrices. It offers the user the possibility to set more specific parameters, such as the number of mismatches allowed in every individual pattern element, and gain statistical information on the mapping output. These quality metrics can help to debug novel sequencing technologies, especially in the developmental phase.

Tools like kITE and alevin are specialized for processing genomic data. These tools focus on the extremely fast and memory efficient mapping of transcripts to a reference genome and added features to handle existing single-cell protocols. In contrast, we developed ESGI with a focus on more flexibility in the demultiplexing step, making it especially suitable for novel experimental designs with complex barcoding schemes. ESGI is developed as a multi-threaded, RAM efficient tool. However, it relies on STAR for mapping of genomic sequences and therefore lacks the efficiency of software specialized for pseudoalignment, such as kallisto or alevin.

ESGI is primarily designed for read demultiplexing and cell assignment, and therefore relies on STAR as an alignment backend to obtain genomic coordinates and gene annotations. In this setup, STAR is effectively used in an exon-centric manner, as plain STAR gene assignment is restricted to exonic regions. Pipelines such as zUMIs or Cell Ranger integrate additional steps to explicitly support intronic counting (for example, via featureCounts [33] or internal intron-aware logic). Nevertheless, ESGI can be used to perform demultiplexing with flexible barcode pattern definitions and indel-aware barcode matching, after which the resulting RNA reads can be processed with downstream tools that support alignment to intronic as well as exonic regions.

Allowing for insertions and deletions (indels) may have limited impact in simple barcoding schemes with high sequence quality and saturated sequencing depth. However, explicit handling of indels becomes increasingly important for lower-quality data and for technologies employing complex barcoding strategies. The prevalence of indels is influenced by multiple factors, including barcode and sequence design, experimental protocol, and the number of amplification and barcoding steps. Especially in combinatorial barcoding approaches involving multiple rounds of barcode attachment, indels can shift the positions of downstream barcodes. ESGI explicitly addresses this challenge by implementing indel-aware barcode matching within user-defined flexible barcode patterns and can accurately identify barcodes even when their positions are altered.

In conclusion, ESGI supports processing of flexible barcode designs by enabling generic barcode patterns, indel-aware barcode processing and variable-length barcodes. These properties make ESGI particularly well-suited for emerging sequencing technologies, for which standardized processing tools may not yet be available. We hope that ESGI becomes a useful tool for processing novel multiplexed single-cell sequencing technologies, without the need for tweaking existing tools or creating custom workflows. Additionally, the statistical information on the quality of mapping that ESGI provides, such as errors at various barcode positions, can provide valuable insight for refining highly multiplexed new technologies during their early development.

## CODE AVAILABILITY

ESGI is written in C++ and released under GNU General Public License (GPL) v3.0. ESGI can be obtained either as ready-to-use precompiled binaries or by compiling the source code, both of which are freely available at https://github.com/tstohn/ESGI. A documentation with examples how to use ESGI is available at: https://tstohn.github.io/ESGI.documentation. The code for the analyses of this paper is available at https://github.com/tstohn/Analysis-ESGI.

## A SUPPLEMENTARY FILES

### A.1 Multi-pattern mapping

Hierarchical and dependency-based barcode patterns pose a particular challenge for conventional demultiplexing tools, as they violate the assumption of fixed-length, positionally independent barcode elements. In such designs, the identity and length of barcode segments may depend on preceding elements, leading to variable-length and structurally interdependent barcode patterns. One example is scID-seq, a well-based technology to measure single-cell protein abundances [2]. The technology employs a stagger-based design in which the set of valid barcodes within a barcode element is constrained by the length of an upstream stagger element (Fig. S1a). ESGI’s demultiplexing module enables explicit modeling of inter-element dependencies by representing each valid stagger–barcode combination as an independent pattern (Fig. S1b). For each read, ESGI systematically evaluates all specified patterns and assigns barcodes when a unique pattern match is identified.

**Figure S1:**
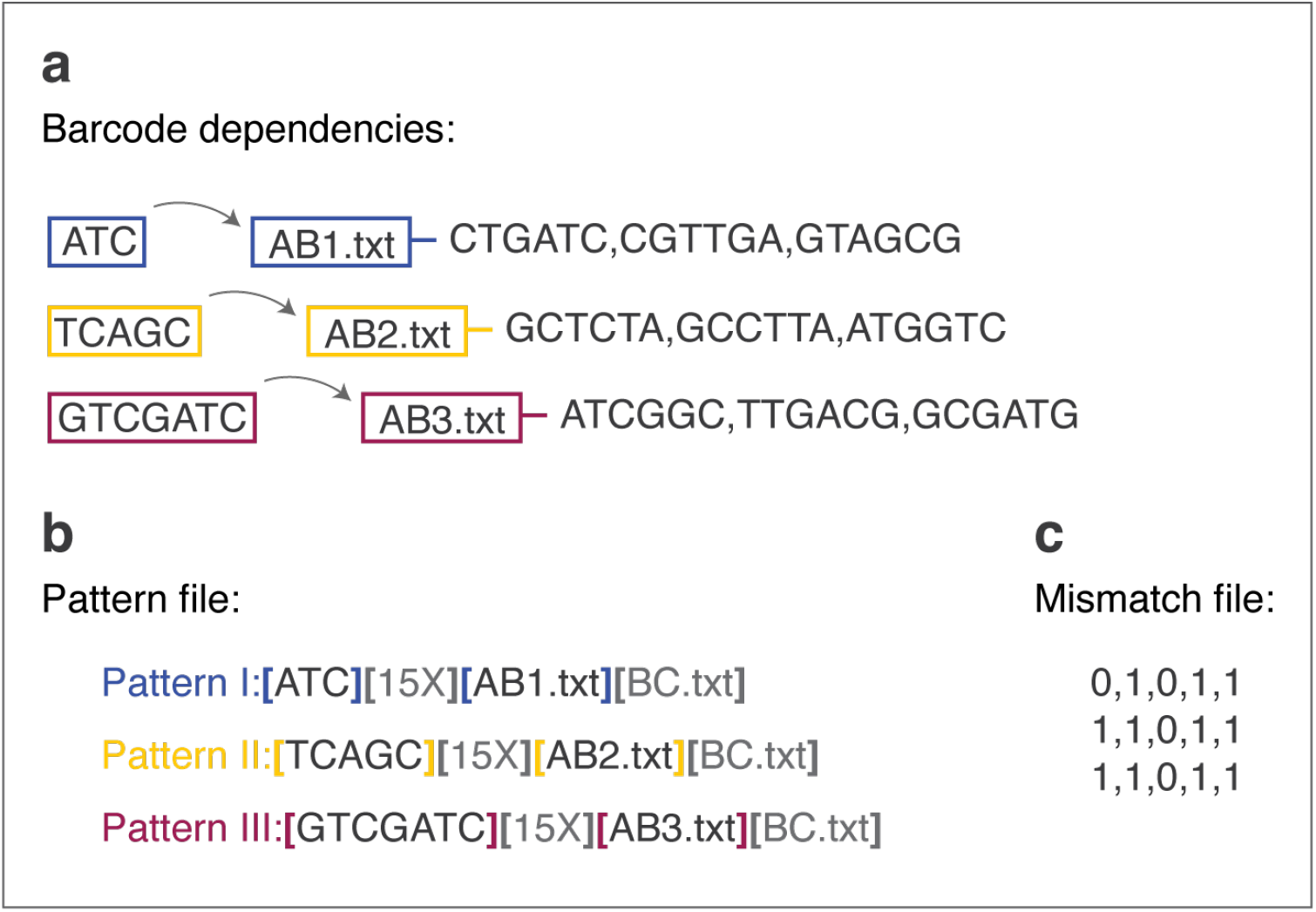
Demultiplexing multipattern designs with barcode dependencies. A.) To expand the barcode design space, dependencies can be introduced between discrete barcode elements within a pattern. In this example, the identity encoded by a barcode element is constrained by the length of a constant stagger element (ATC, TCAGC, GTCGATC). These staggers vary in length, and each is associated with a specific substring of potential sequences for the barcode element. B.) Each unique stagger-barcode combination is assigned a distinct pattern name and set of pattern elements. Each element is enclosed in brackets and can contain a constant sequence (stagger), random sequence (15X), or barcode (.txt). Pattern-specific files (AB.txt) encode the feature identity, while shared files (BC.txt) encode the single-cell identity. C.) For each pattern, the maximum allowed mismatches per element are defined as a comma-separated list of integers.

### A.2 Barcode mapping procedure in demultiplex

`demultiplex` identifies barcode patterns in sequencing reads by matching the user-defined pattern to input sequences (e.g., FASTQ reads). An overview of the mapping strategy is shown in Supplementary Fig. S2.

#### A.2.1 Hash-based barcode lookup

To efficiently resolve barcode identities at positions where multiple barcode candidates are possible, `demultiplex` uses as a first step precalculated hash tables (Supplementary Fig. S2b). It performs direct lookups using a precomputed substitution map that links every possible sequence (including sequences containing a single substitution) to the corresponding original barcode. Per default a second map including all sequences with a single insertion or deletion (Levenshtein distance of one) is used. This indel-aware map can be disabled to reduce memory usage when only Hamming-distance matching is required. Only uniquely mapped reads are retained.

#### A.2.2 Positional *k*-mer filtering

To keep memory low possible sequences with more than two mismatches are not precalculated. When more than one mismatch is allowed, or when indel-aware matching is requested without using the precomputed Levenshtein map, `demultiplex` applies a positional *k*-mer filtering strategy (Supplementary Fig. S2b).

For all candidate barcodes, the algorithm generates every possible *k*-mer and stores it together with its positional index as a hash key. Each key maps to the barcode(s) in which the corresponding positional *k*-mer occurs. During filtering, positional *k*-mers are generated from the observed sequence and matched against this map to retain only compatible barcode candidates. A barcode is retained if it shares at least a minimum number of positional *k*-mers with the observed sequence. The minimum number of required matches is defined as:

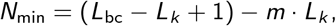

where *L*_bc_ denotes the barcode length, *L*_*k*_ the *k*-mer length, and *m* the maximum number of allowed mismatches.

To account for indels, positional indices of *k*-mers are allowed to shift within a window that is of length of the number of permitted mismatches (positional *k*-mers can occur at the expected position +/-mismatches).

#### A.2.3 Final alignment

After filtering, remaining candidate barcodes are aligned to the extracted read segment. If a unique best match is identified, the read is assigned accordingly. Only uniquely mapped reads are retained. Barcodes are mapped with a semi-global alignment strategy (Fig. S3).

### A.3 UMI collapsing

Various UMI collapsing strategies have been proposed and how UMIs are collapsed has a strong influence on the number of counts that are detected (Fig. S4). Collapsing UMIs wrongfully leads to an under or overestimation of counts, especially for features that are generally highly abundant. Many pipelines like zUMIs [14], Cell Ranger [12] or alevin [15] consider mismatches when collapsing UMIs, while other tools like kallisto and bustools [17] collapses UMIs without allowing for mismatches, which can lead to over-collapsing of UMIs. Many methods apply graph-based approaches merging similar UMIs like in alevin. The tool dropEst provides a Bayesian approach to demultiplex UMIs [34]. During UMI collapsing some tools additionally consider the base quality or abundance of reads to inform the collapsing of UMIs. Alevin for example collapses the less abundant UMI only into the more abundant one if its counts are less or equal to half the counts of the more abundant UMI (equivalent to the 50% collapsing rule in (Fig. S4)). A benchmark of various processing strategies for scRNA-seq data with UMI counts found that various UMI preprocessing methods had less of an impact on the results of the analysis than for example normalization strategies [35]. Nevertheless, for optimal results the collapsing of UMIs should be handled with care as counts can differ a lot depending on the chosen UMI collapsing strategy (Fig. S4).

To enable user-defined UMI collapsing, we provide two parameters: mismatches (default = 1), which specifies the maximum allowed Hamming distance between UMIs, and umiAbundanceThreshold (default = 0.2), which defines the relative abundance threshold for error correction. UMIs within the specified Hamming distance are eligible for collapsing if the less abundant UMI has fewer counts than the specified fraction (between 0 and 1) of the more abundant UMI. By default, UMIs differing by one mismatch are collapsed when the lower-abundance UMI has less than 20% of the counts of the higherabundance UMI.

**Figure S2:**
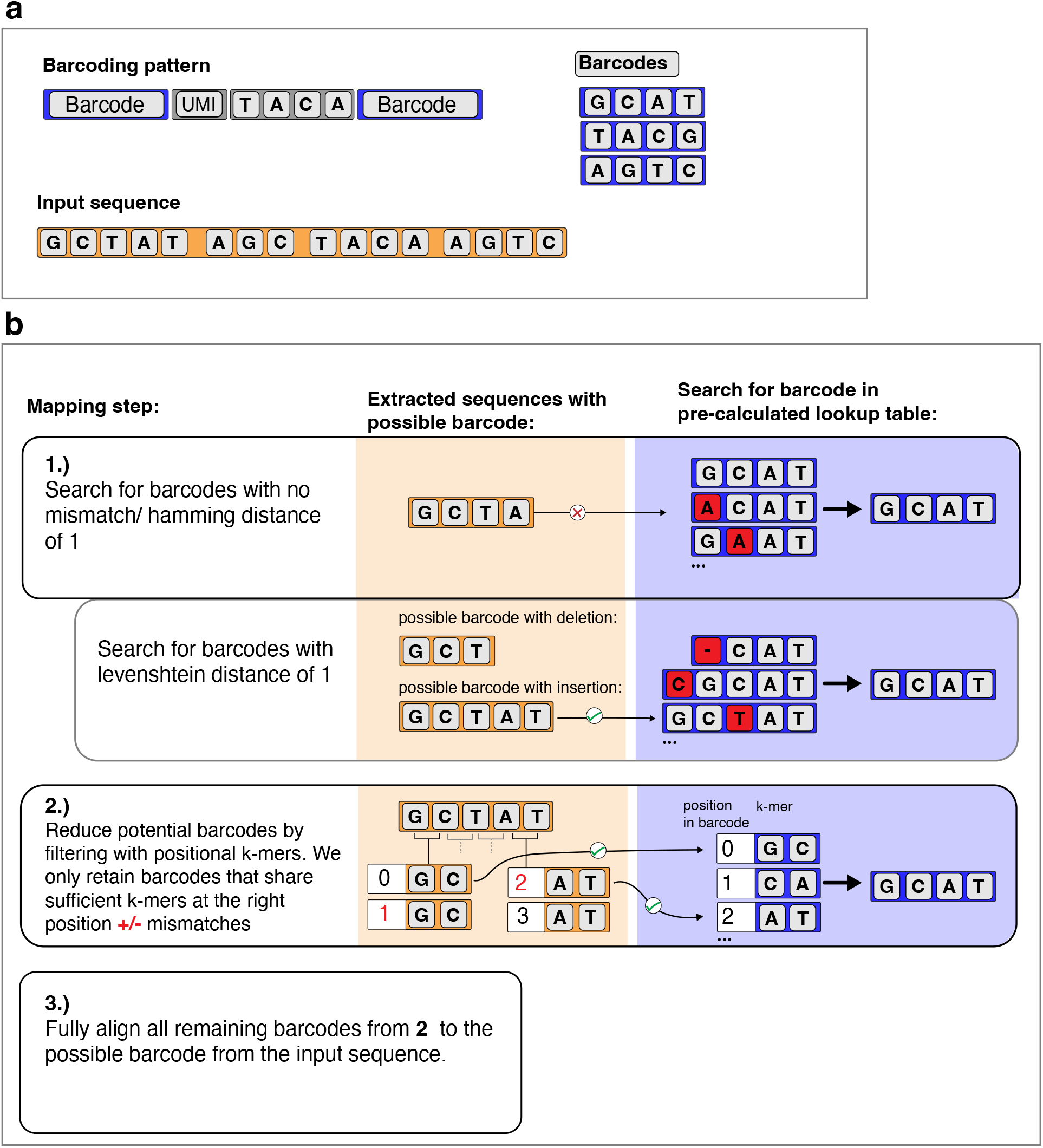
Overview of the barcode mapping workflow. (a) Example barcode pattern consisting of multiple barcode elements (blue), a UMI and a constant sequence (grey) that are mapped to input reads (yellow). (b) Three-stage barcode identification strategy combining hash-based direct lookup and positional *k*-mer filtering, followed by final alignment.

**Figure S3:**
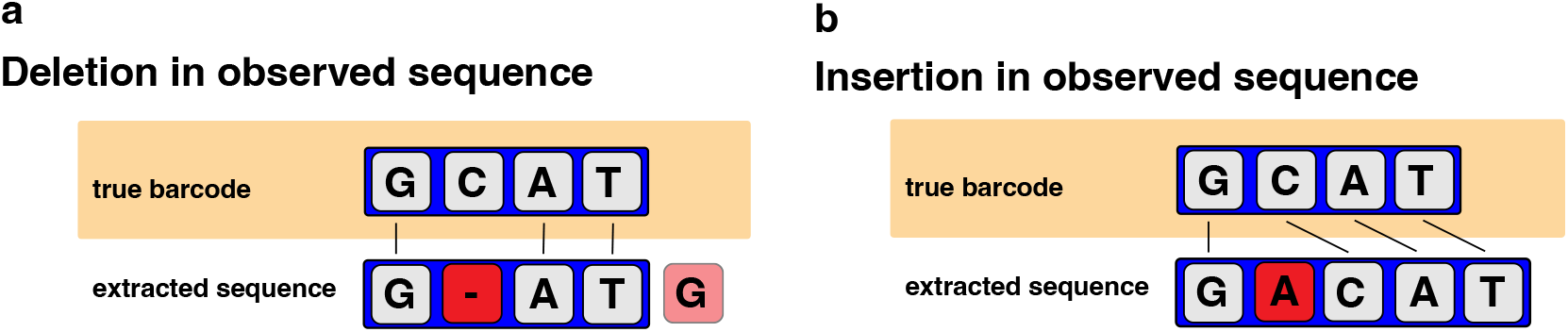
Semi-global alignment for barcode demultiplexing. Let *L* denote the expected barcode length and *k* the maximum number of allowed mismatches, including insertions and deletions. demultiplex extracts a sequence of length *L* + *k* from a FASTQ read and aligns it to the expected barcodes using semi-global alignments. In this example, *L* = 4 and *k* = 1, such that a sequence of length 5 is extracted. (a) When the number of deletions exceeds the number of insertions, the observed sequence contains additional bases that are not part of the mapped barcode (blue background). The semi-global alignment allows unpenalized overhangs at the end of the mapped barcode in the extracted sequence (light red), while deletions occurring within the barcode are penalized (dark red). (b) When the number of insertions does not exceed *k*, the observed sequence has the correct effective length, enabling correct barcode identification within the whole length of the extracted sequence.

**Figure S4:**
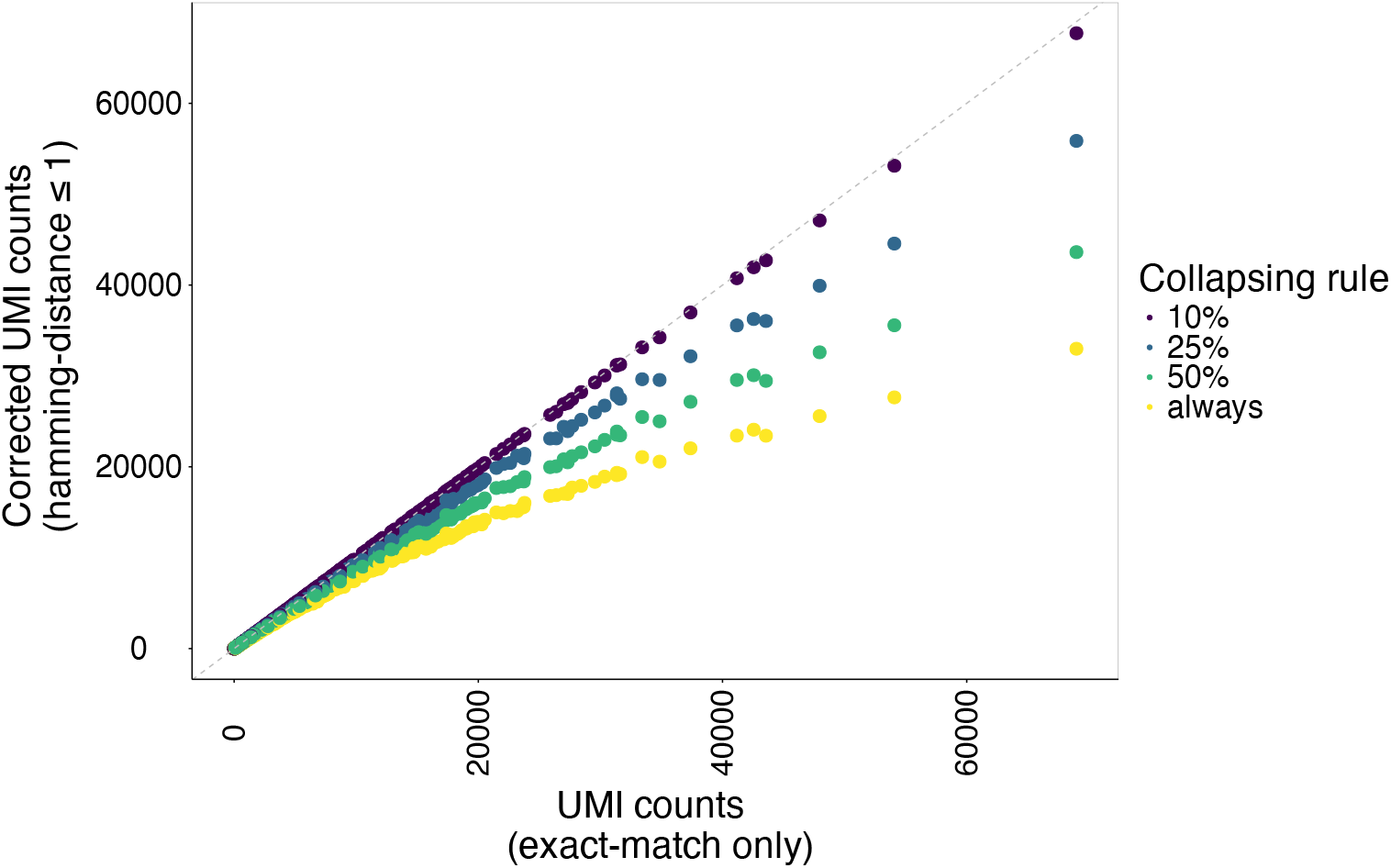
Collapsing of UMIs with ESGI (count-tool) for the protein modality of the SIGNAL-seq dataset. For every feature in every detected cell collapsed UMI counts without error correction (0 mismatches) are plotted against UMI counts that were corrected for mismatches (Hamming distance *<*= 1). Additionally, UMIs were only collapsed if the less abundant UMI was less than 10%, 25%, 50% of the more abundant UMI by setting the umiAbundanceThreshold parameter accordingly, or it was collapsed regardless of the UMI abundance (always).

**Figure S5:**
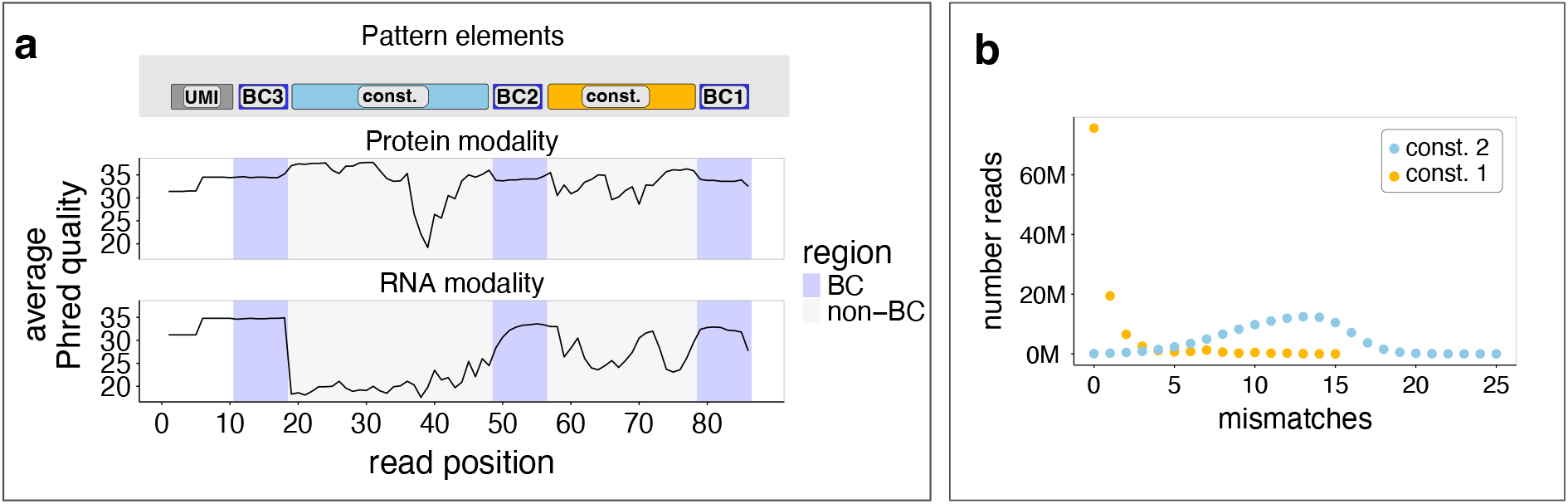
Average base quality of the reverse read containing the three single cell barcodes of the SIGNAL-seq data. (a) The average Phred score per position for the bases in the FASTQ file of the protein and the RNA modality. Because ESGI allows the mismatch tolerance to be defined separately for each pattern element, users can adapt the permitted number of mismatches according to the observed sequencing quality (e.g., Phred score distributions) of individual barcode or constant regions. (b) The number of mismatches in the two constant elements in the pattern for the RNA modality. We ran ESGI allowing for 15 mismatches in the first constant region and 25 mismatches in the second constant region and counted the number of mismatches (up to 15 and 25) for all reads that could be successfully mapped to the pattern.

**Figure S6:**
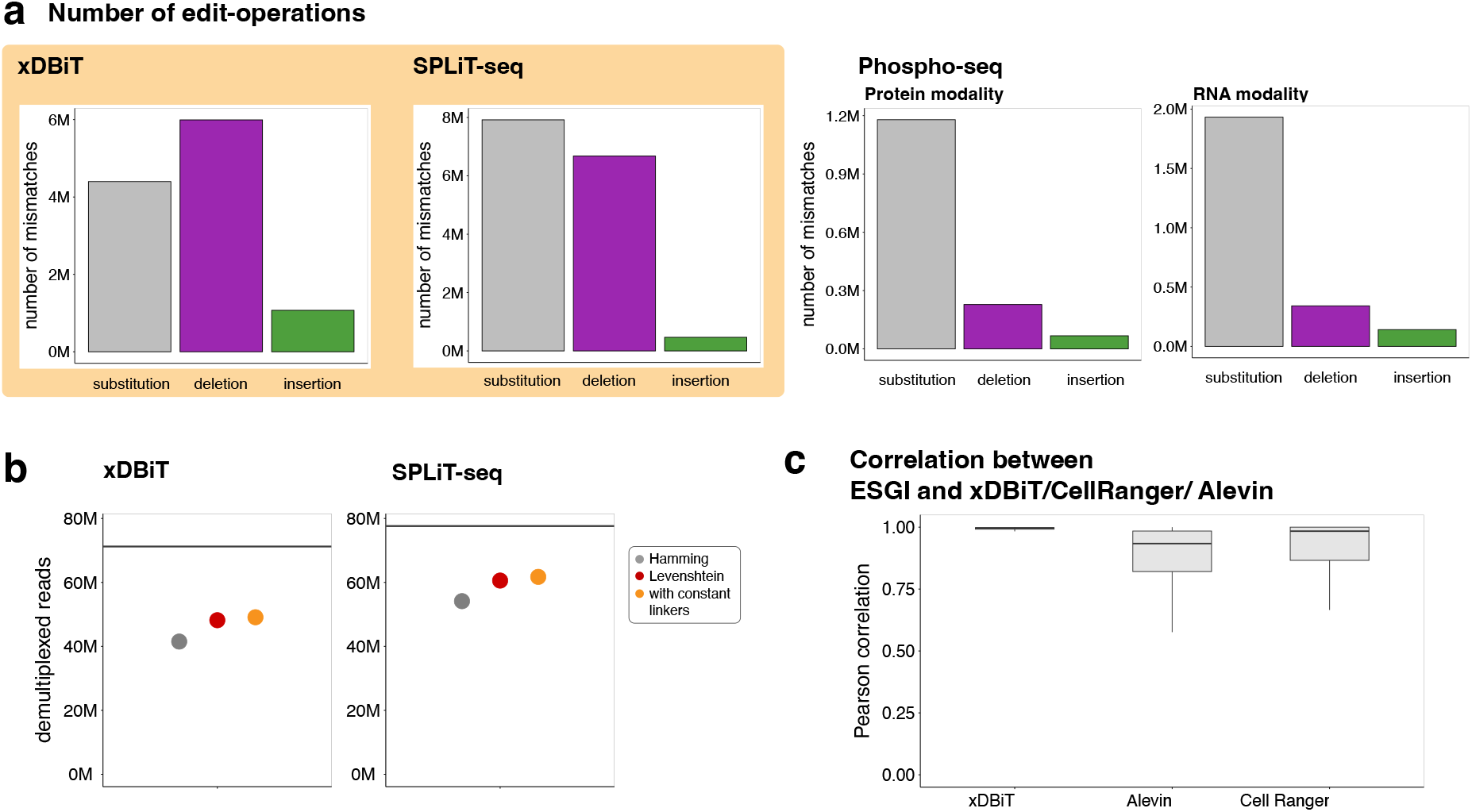
Indel-aware demultiplexing of the xDBiT, SPLiT-seq and Phospho-seq (protein and RNA) datasets. (a) Number of edit-operations in the barcode elements of the datasets. Datasets with several barcode elements encoding single-cell indices are highlighted in yellow. (b) Successfully demultiplexed reads using a Hamming distance, a Levenshtein distance, or a Levenshtein distance in combination with mapping the constant linker sequences between barcode elements explicitly. The horizontal lines represent the total number of reads in the raw FASTQ files. (c) Pearson correlations of feature counts (protein or RNA counts) of the indicated tool with ESGI per cell after UMI collapsing.

**Figure S7:**
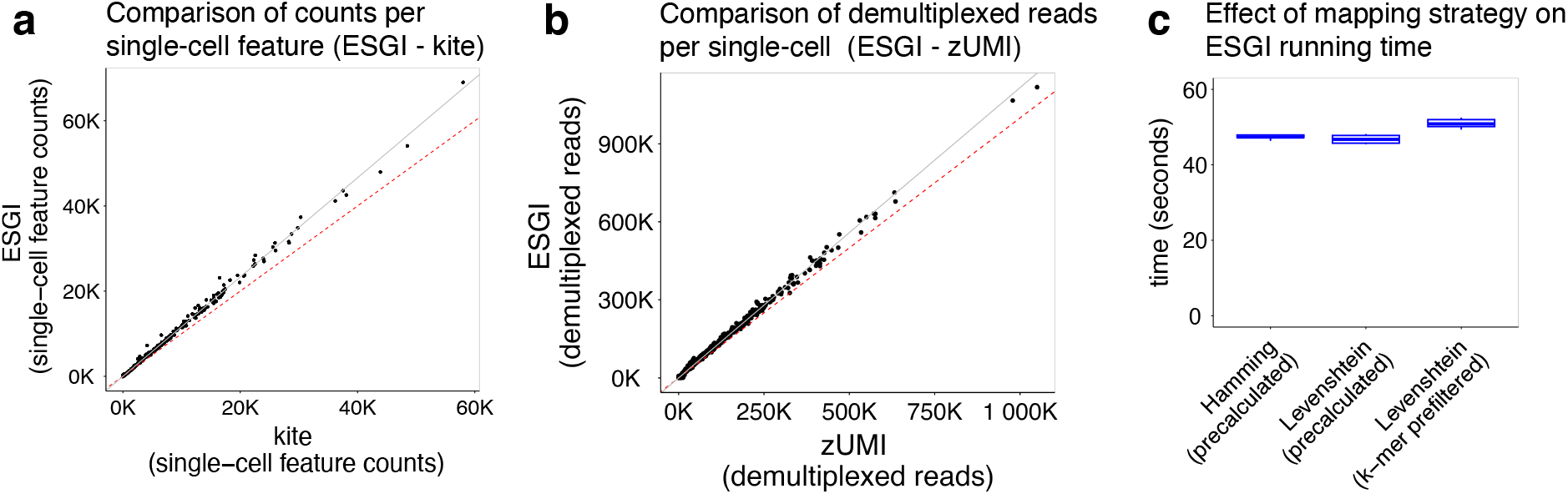
Indel aware demultiplexing using ESGI results in more counts per feature without increasing runtime. (a) Scatterplot comparing the counts per feature per cell when demultiplexing the protein modality of SIGNAL-seq data with either ESGI or kITE, when allowing for one mismatch in the barcode elements and collapsing UMIs without error correction (default of kITE). Dashed red line indicates equal counts. (b) Scatterplot comparing the number of demultiplexed reads for zUMIs and ESGI of the RNA modality of the SIGNAL-seq data. Counting genomic sequences with UMI correction is more complex than counting feature barcodes. zUMIs and ESGI differ in multiple processing steps, making direct comparison of count matrices challenging. In particular, differences in UMI collapsing rules and genome alignment and gene assignment strategies can substantially influence the resulting counts. Therefore, we compare the number of reads that could be demultiplexed before counting features in single-cells. (c) Running time of ESGI on the protein modality of the SIGNAL-seq dataset was evaluated under three demultiplexing strategies: (i) precalculation of barcodes within a Hamming distance of one; (ii) precalculation of barcodes within a Levenshtein distance of one; and (iii) precalculation restricted to Hamming distance of one while allowing indels through direct barcode-to-sequence alignment, with the candidate barcode set reduced using positional *k*-mers (see Figure S2).

